# CO_2_-concentrating nanoparticles drive progressive photosynthetic and cell-wall remodelling in *Arabidopsis thaliana* cells

**DOI:** 10.64898/2026.10.30.763161

**Authors:** Cyril Routier, Denise Pagliara, Anna Karlsson, Lucija Lisica, Léo Boutonnet, Léa Bogdziewiez, Erica Colaprico, Ariel García Fleitas, Xu Jin, Åsa Strand, Daniele Mantione, Stéphane Verger, Elton P. Hudson, Eleni Stavrinidou

**Affiliations:** Laboratory of Organic Electronics, Department of Science and Technology, Linköping University, SE-60174, Norrköping, Sweden; School of Engineering Sciences in Chemistry, Biotechnology and Health, Science for Life Laboratory, KTH - Royal Institute of Technology; Stockholm, Sweden; Umeå Plant Science Centre, Department of Forest Genetics and Plant Physiology, Swedish University of Agricultural Sciences, SE-90183 Umeå, Sweden; Umeå Plant Science Centre, Department of Plant Physiology, Umeå University, SE 901-87 Umeå, Sweden; POLYMAT, University of the Basque Country UPV/EHU, 20018 San Sebastián, Spain; Ikerbasque, Basque Foundation for Science, Bilbao, Spain; Wallenberg Initiative Materials Science for Sustainability, Linköping University, SE-60174, Norrköping, Sweden

**Author notes:** equal contribution (Cyril Routier, Denise Pagliara).

**Keywords:** plant cell biohybrids, CO_2_ capture, polyethyleneimine nanoparticles, photosynthesis, cell wall biosynthesis

## Abstract

Plant cells offer a distinctive platform for biohybrid engineering because they combine growth and proliferation with photosynthetic carbon acquisition and the continuous fabrication of structurally complex biomass. Harnessing these capabilities could enable living systems in which synthetic functionality develops together with cellular physiology and material organization. Yet nanomaterial engineering of proliferating plant cell populations remains largely unexplored. Here, we establish a biohybrid system combining photosynthetically active *Arabidopsis thaliana* suspension cells with chitosan-modified polyethyleneimine (gPEI–Chi) nanoparticles, a synthetic CO_2_-concentrating nanomaterial. By supplying fresh nanoparticles at each subculture, newly generated cell populations were successively functionalized, driving progressive changes in cellular and multicellular phenotypes. Nanoparticle functionalization reduced net headspace CO_2_ accumulation, increased chlorophyll content and induced stage-dependent reorganization of photosynthetic function and carbon-handling pathways. These physiological changes were accompanied by altered cell wall regeneration and progressive remodelling of cell wall composition, spatial organization and ultrastructure, ultimately promoting more cohesive multicellular aggregates. Our findings establish successive nanomaterial functionalization as a strategy to steer the evolving physiology and structural organization of proliferating plant cell populations, opening opportunities for plant biohybrid systems and engineered living materials.

## Introduction

Living cells offer capabilities difficult to reproduce in conventional engineered systems as they grow, multiply, sense their environment and continuously remodel their own structures. Integrating synthetic functionality into proliferating cell populations could therefore enable biohybrid systems that develop as their living components grow. Plant cells are particularly attractive because they combine photosynthetic carbon acquisition with the ability to convert assimilated carbon into biomass and structurally complex cell walls^1^, providing an opportunity to engineer both cellular physiology and biological material formation.

Plant nanobionics integrates engineered nanomaterials with living plants to introduce or enhance biological functions through nano–bio interactions^2,3^. Nanomaterials have enabled molecular delivery^4,5^, sensing^6,7^ and modulation of photosynthetic processes^8,9^ despite the transport barriers imposed by plant cell walls and membranes^10,11^. These approaches have largely focused on intact plants and differentiated tissues, whereas proliferating plant cells remain comparatively unexplored. Suspension cells continuously divide and can be propagated through successive subcultures, creating an opportunity to functionalize newly generated cell populations and investigate whether nanomaterial interactions can progressively reshape cellular and multicellular phenotypes.

Carbon handling provides a particularly relevant target for such engineering. CO_2_ is the principal carbon source for photosynthesis, yet its fixation is constrained by the catalytic properties of RuBisCO, including competing oxygenase activity, and by limitations in CO_2_ transport to the site of carboxylation^12,13^. Natural carbon-concentrating mechanisms increase CO_2_ availability around RuBisCO and suppress photorespiration^14,15^, while efforts to engineer such mechanisms in other photosynthetic organisms have largely relied on genetic and metabolic modification^16–18^. Synthetic materials capable of concentrating CO_2_ offer an alternative strategy to modify the carbon environment of photosynthetic cells without editing their genome. Although nanomaterials have been used to modulate light harvesting^19,20^ and photosynthetic electron transport^8,21^, comparatively little attention has been directed towards engineering inorganic carbon availability^22^.

Building on the ability of polyamines to capture CO_2_^23–25^, we previously developed green-fluorescent chitosan-modified polyethyleneimine nanoparticles (gPEI–Chi NPs) as a synthetic CO_2_-concentrating nanomaterial for plants^26^. Combining the CO_2_ affinity of polyethyleneimine with the phytocompatibility of chitosan^27,28^, gPEI–Chi NPs reversibly capture inorganic carbon and increased RuBisCO carboxylation by 20% in enzymatic assays. Following infiltration in planta, the nanoparticles (NPs) distributed throughout tobacco leaf tissues, including chloroplast-rich regions, while maintaining phytocompatibility^29^. These findings established the feasibility of manipulating the local inorganic carbon environment of photosynthetic tissues using nanomaterials. However, how such functionality interacts with proliferating plant cells as they divide and are successively propagated, and whether functionalization of newly generated cell populations can progressively reshape their phenotype, remain unknown.

This question extends beyond photosynthesis because assimilated carbon also provides the building blocks for the polysaccharide-rich plant cell wall. Changes in carbon handling could therefore influence both cellular metabolism and the structural organization of multicellular assemblies. This coupling is particularly relevant to engineered living materials (ELMs), where living cells provide adaptive and regenerative functionality^30,31^. Plant cells are attractive building blocks for such systems because they photosynthetically acquire carbon and convert it into structurally complex cell walls. Recently, we demonstrated that plant cell-based ELMs can exploit cellular metabolic plasticity and cell wall biochemistry to generate adaptive and programmable material properties^32^. Engineering proliferating plant cells with nanomaterials could therefore provide a route to regulate both cellular physiology and structural organization as living populations develop.

Here, we establish a biohybrid system combining photosynthetically active *Arabidopsis thaliana* suspension cells with CO_2_-concentrating gPEI–Chi NPs and investigate how NP functionalization across successive subcultures reshapes cellular and multicellular phenotypes. We show that successive functionalization alters carbon handling and photosynthetic function while progressively remodelling cell wall regeneration, composition and organization, ultimately promoting a more cohesive multicellular phenotype. These findings demonstrate that nanomaterial functionalization of proliferating plant cell populations can couple changes in cellular physiology with structural reorganization, providing a platform for plant biohybrid systems and engineered living materials.

### gPEI–Chi NPs functionalize photosynthetic Arabidopsis suspension cells

*Arabidopsis thaliana* suspension cultures were established from root-derived calli and adapted to photomixotrophic growth under constant illumination and reduced sucrose supplementation (Fig. 1a), as previously demonstrated^33^. These conditions promoted chloroplast development while maintaining suspension growth, with confocal imaging revealing robust chlorophyll autofluorescence throughout the cultures (Fig. 1b).

**Fig. 1.**
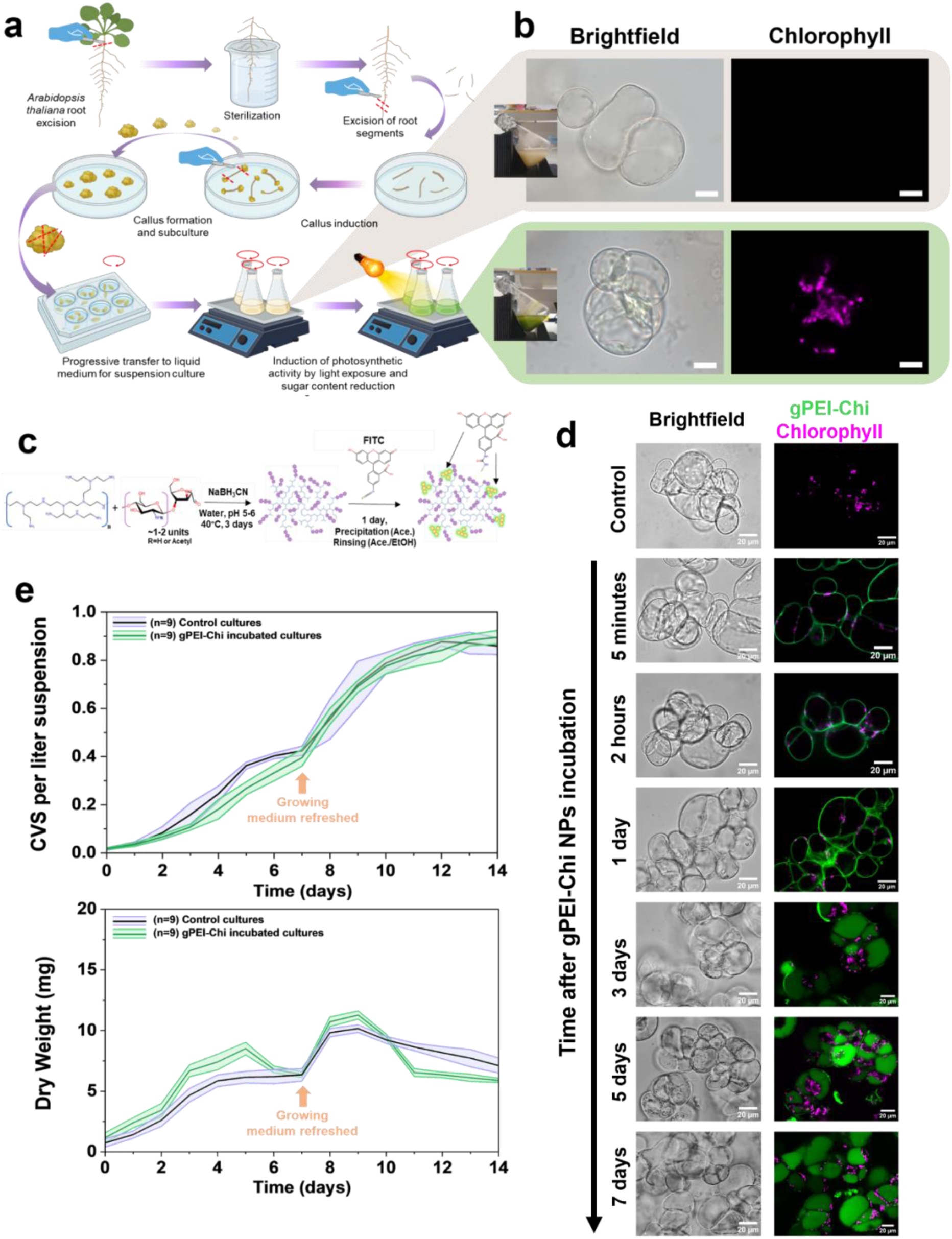
gPEI-Chi NPs are internalized by photosynthetic *Arabidopsis thaliana* cells in suspension cultures without inducing cytotoxic effects. **a**, Schematic representation of the generation of photosynthetic suspension cultures derived from *Arabidopsis thaliana* root explants (created in BioRender.com). **b,** Confocal microscopy images of cells showing brightfield and chlorophyll autofluorescence, confirming the establishment of photosynthetically active cells. Scale bar 10µm. **c**, Synthesis pathway of green-fluorescent chitosan-modified polyethyleneimine nanoparticles (gPEI-Chi NPs). **d,** Time-course confocal imaging of NP internalization showing progressive association with the cell wall and intracellular localization, with significant uptake observed after approximately 3 days. Scale bar 20µm. **e**, Growth kinetics of control and gPEI-Chi-treated cultures over 14 days, assessed by cell volume sedimentation (CVS, i) and dry weight measurements (ii), demonstrating no significant impact of NP treatment on cell proliferation.

Green-fluorescent chitosan-modified polyethyleneimine nanoparticles (gPEI–Chi NPs), previously developed as CO_2_-capturing nanomaterials for in planta applications^26^, were synthesized by reductive amination of polyethyleneimine with oligochitosan followed by fluorescein isothiocyanate (FITC) labelling (Fig. 1c). NPs were added at subculture at a final concentration of 0.1 mg mL⁻¹. Evans Blue staining revealed no detectable increase in cell death following NP treatment in either non-photosynthetic or photosynthetic cultures (Supplementary Fig. S1). Notably, membrane-compromised cells lacked detectable NP fluorescence, suggesting that intracellular NP accumulation depends on cellular integrity.

Upon addition, gPEI–Chi NPs rapidly associated with the cell periphery, consistent with PEI affinity for plant cell wall components (Supplementary Fig. S2)^26,34,35^. NP-associated fluorescence subsequently redistributed towards intracellular compartments, with substantial internalization observed after approximately 3 days (Fig. 1d). Spectroscopic analysis supported progressive cellular association and uptake as FITC-associated absorbance and fluorescence in NP-containing medium decreased substantially faster in the presence of cells, approaching baseline by day 5 (Supplementary Fig. S3). Consistently, confocal quantification showed increasing NP-associated fluorescence within the cellular area and progressive redistribution from the cell surface towards intracellular compartments over 6–7 days (Supplementary Fig. S4). Together, these measurements demonstrate progressive NP uptake and intracellular accumulation during culture.

To determine whether NP functionalization affected culture growth, cell volume after sedimentation (CVS; Fig. 1e-i and Supplementary Fig. S5) and biomass dry weight (Fig. 1e-ii) were quantified over 14 days. NP treatment did not substantially alter the characteristic growth kinetics of either photosynthetic or non-photosynthetic cultures, although a modest transient delay was observed during early growth (Fig. 1e-i and Supplementary Fig. S6a). Photosynthetic cultures required medium renewal after 7 days, when NPs were replenished at the same concentration.

Despite similar CVS profiles, NP-treated photosynthetic cultures showed transient changes in dry biomass. Dry weight was 43% and 38% higher than controls at days 3 and 5, respectively, before returning to control levels by days 6–7. Following medium renewal, biomass increased again by 9–11% at days 8–9, decreased by approximately 24% at day 11 and subsequently stabilized near control levels. The divergence between CVS and dry weight indicates that NP functionalization transiently alters biomass accumulation and/or cellular composition without substantially affecting culture proliferation.

Together, these results demonstrate that gPEI–Chi NPs efficiently associate with and are internalized by photosynthetic *Arabidopsis thaliana* suspension cells while preserving cellular viability and proliferative capacity.

### gPEI–Chi NP functionalization across successive subcultures induces progressive phenotypic changes

Despite efficient intracellular association, a single NP functionalization cycle produced no major macroscopic phenotypic changes beyond transient differences in biomass accumulation. Because cellular responses to sustained abiotic stimuli can develop beyond acute responses to encompass longer-term metabolic, structural and population-level changes, we investigated whether repeated gPEI–Chi NP functionalization across successive subcultures would progressively alter the phenotype of photosynthetic cultures^36–38^.

Photosynthetically active *Arabidopsis thaliana* suspension cultures were subcultured weekly with fresh NPs supplied at the same concentration at each subculture to maintain exposure as the cells proliferated (Fig. 2a). Gen0 denotes the initial functionalization whereas Gen3 and Gen6 represent cultures maintained under repeated NP functionalization for three and six successive weekly subcultures, respectively.

**Fig. 2.**
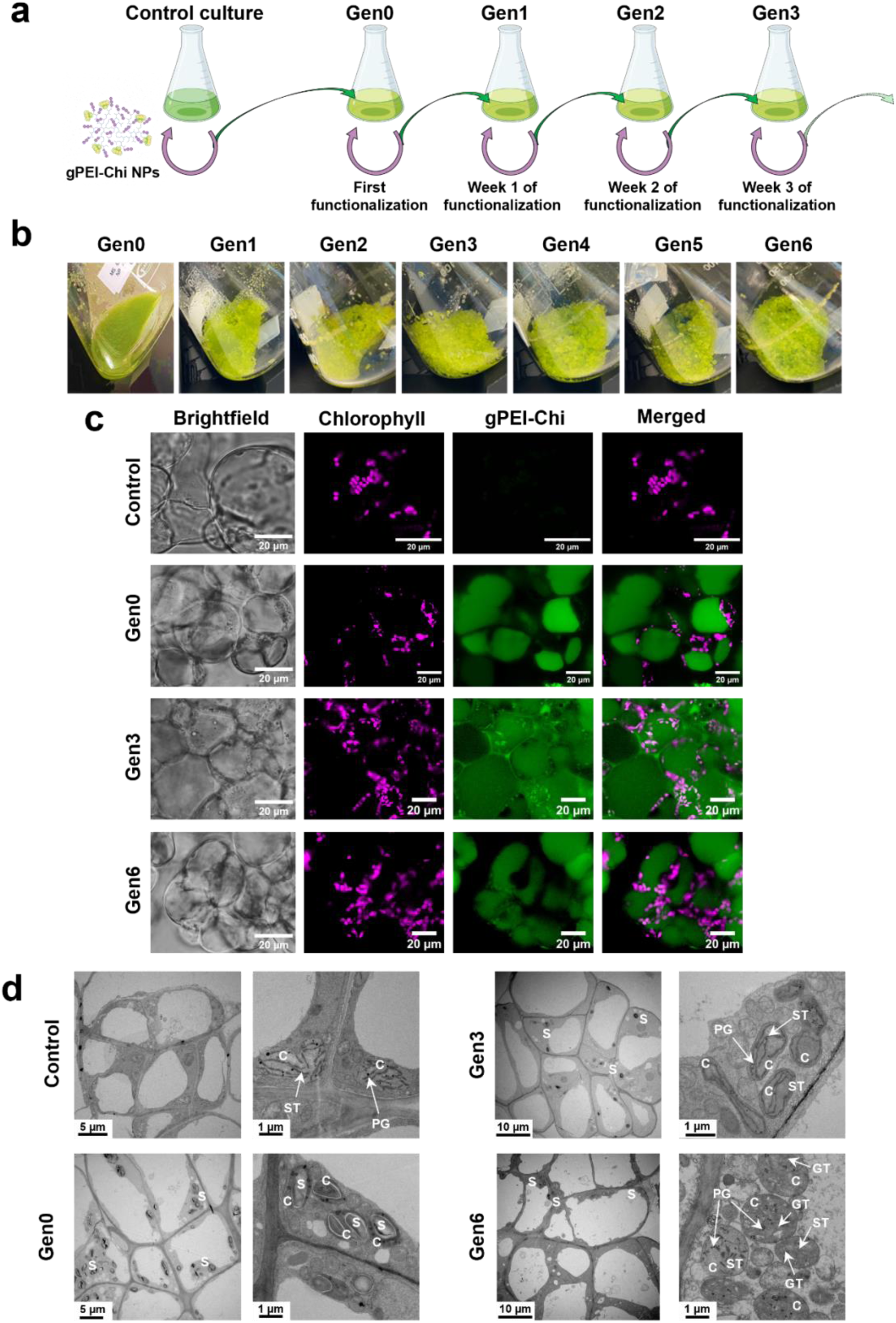
gPEI-Chi NPs functionalization over generations of cells induces cellular aggregation and alters cell morphology and chloroplast development. **a**, Schematic representation of the experimental workflow for multi-generational NP functionalization, in which suspension cultures are repeatedly exposed to gPEI-Chi NPs across successive subcultures (Gen1, Gen2, etc.). **b**, Representative pictures showing the progressive macroscopic appearance of cultures over generations, highlighting increased aggregation and changes in culture morphology upon repeated NP exposure. **c**, Confocal microscopy images of NP-treated cultures (Gen6) revealing enhanced cellular aggregation and increased chlorophyll autofluorescence, indicative of more developed chloroplast structures, colocalized with gPEI-Chi NP fluorescence in the case of Gen3 and Gen6 cultures. **d,** Transmission electron microscopy (TEM) images of NP treated and non-treated cultures, underlying the ultrastructural differences between the conditions. Control cultures present chloroplasts with abundance of stromal thylakoids and plastoglobuli (black dots). Gen0, Gen3 and Gen6 present high content of starch-enriched chloroplasts and their number increases particularly in Gen3 and Gen6. In Gen6, granal thylakoids can be also detected together with stromal thylakoids. In the images C=chloroplasts, S=starch, ST=stromal thylakoids, PG=plastoglobuli, GT=granal thylakoids.

Over successive subcultures, NP-functionalized cultures progressively developed a distinct macroscopic phenotype characterized by denser, more cohesive cell clusters relative with control and Gen0 cultures, which remained more homogeneous and exhibited a smoother suspension appearance (Fig. 2b). Confocal imaging confirmed the increased aggregation of Gen3 and Gen6 cultures (Supplementary Fig. S7 and S8). Chlorophyll autofluorescence also appeared more pronounced and broadly distributed in Gen3 and Gen6 cells compared with control and Gen0 cultures, suggesting increased chloroplast development (Fig. 2c). NP-associated fluorescence remained closely associated with chlorophyll-rich intracellular regions in both Gen3 and Gen6 cells, consistent with localization of gPEI-Chi NPs in chloroplast-rich regions. In addition, highly fluorescent shell-like structures were frequently observed in NP-functionalized cultures, consistent with cell wall remnants retaining NP-associated fluorescence after loss of cellular contents (Supplementary Fig. S8b).

Transmission electron microscopy revealed corresponding ultrastructural changes across successive NP functionalization cycles (Fig. 2d, Supplementary Fig. S9). Control cells displayed regular vacuolar and organelle organization, including mitochondria and chloroplasts containing plastoglobules and thylakoid membranes. Gen0 cells exhibited increased aggregation, enlarged vacuoles, more abundant mitochondria and starch-enriched chloroplasts. In Gen3 and Gen6 cultures, cellular aggregation became increasingly pronounced, while vacuoles regained a more regular morphology and chloroplasts appeared more abundant throughout the cytoplasm. Chloroplasts also exhibited increasingly organized thylakoid membranes including grana formation, particularly in Gen6 cells. These observations reveal progressive changes in cellular and chloroplast ultrastructure.

To determine whether successive functionalization altered NP uptake dynamics, Gen3 and Gen6 cultures were monitored following NP replenishment (Supplementary Fig. S10). In both cultures, NP-associated fluorescence rapidly accumulated intracellularly, indicating faster uptake than during the initial Gen0 functionalization. Consistently, FITC-associated absorbance and fluorescence signals appeared in the culture medium immediately after NP replenishment but rapidly declined over subsequent days.

Together, these results demonstrate that gPEI–Chi NP functionalization across successive subcultures induces faster NP uptake and progressively alters culture morphology and cellular ultrastructure resulting in more cohesive cell aggregates and increased chloroplast development.

### NP functionalization reorganizes carbon handling and photosynthetic function

Given the sustained association of gPEI–Chi NPs with chlorophyll-rich intracellular regions, we next investigated whether NP functionalization alters CO_2_ dynamics and photosynthetic function. Net CO_2_ exchange was monitored using a closed-loop system that continuously measured headspace CO_2_ concentrations under isolated conditions (Fig. 3a,b). To assess the intrinsic CO_2_-capturing capacity of the NPs, gPEI–Chi NP suspensions were first monitored in the absence of cells. Headspace CO_2_ remained between 300 and 400 ppm over 7 days, consistently below atmospheric levels, confirming CO_2_ capture by the NPs under the experimental conditions (Supplementary Fig. S11a).

**Fig. 3.**
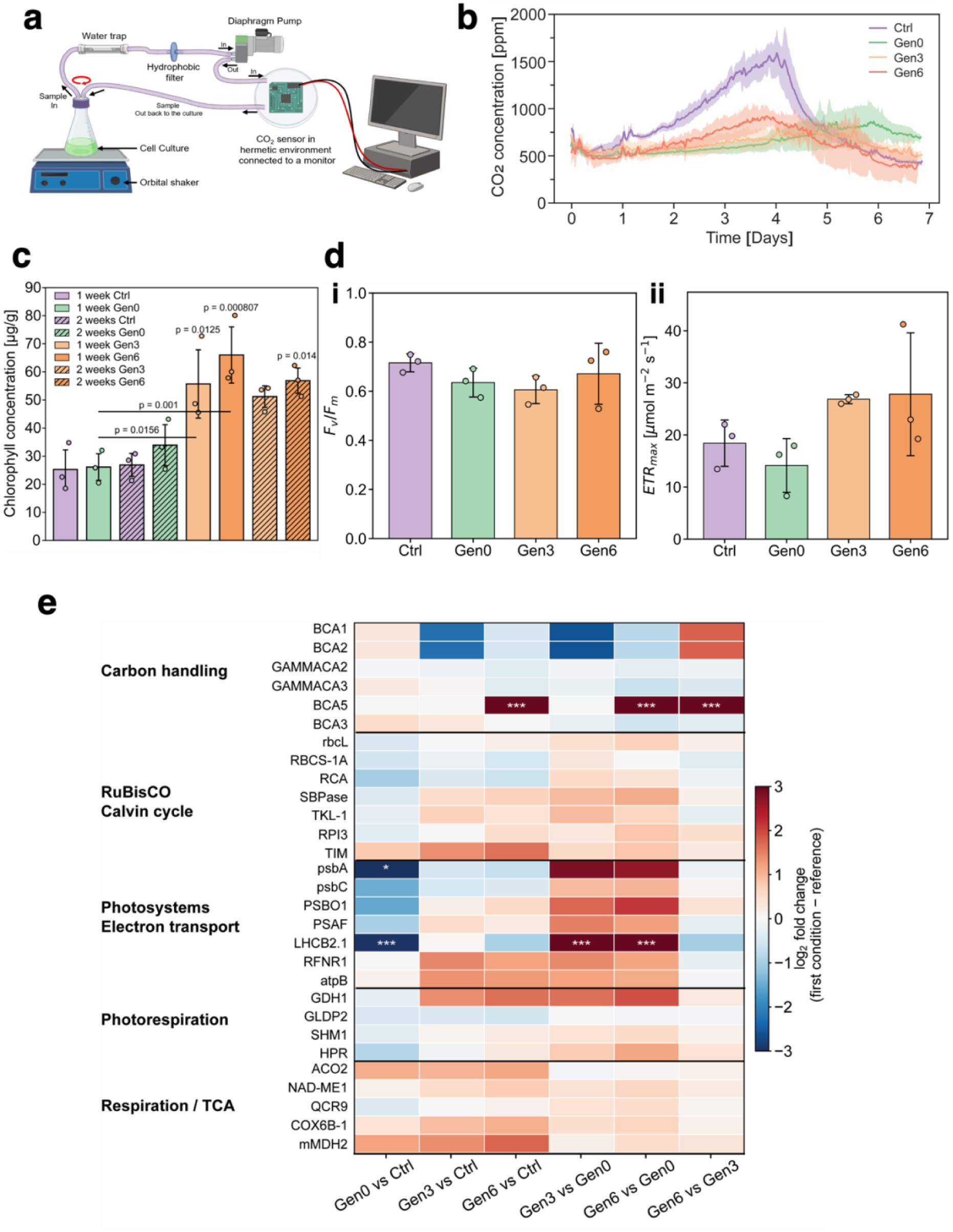
gPEI-Chi NPs enhance CO_2_ capture and modulate photosynthetic activity in plant cell suspension cultures. **a**, Schematic of the closed loop system allowing to measure the CO_2_ concentration in the headspace of sealed culture flasks (created in BioRender.com). **b**, CO_2_ concentration measured in the headspace of sealed culture flasks over 7 days for control (Ctrl), NP-treated (Gen0), and multi-generational cultures (Gen3 and Gen6). NP-treated samples exhibit a greater reduction in headspace CO_2_ compared to controls, while NP-only controls show minimal variation. **c**, Quantification of total chlorophyll content in control and NP-treated cells, showing increased chlorophyll accumulation in functionalized cultures, indicative of enhanced photosynthetic capacity. **d**, (i) Maximum quantum yield of photosystem II (Fv/Fm) and (ii) maximum electron transport rate (ETR_max_) measurement for control cultures and NP-treated cells (Gen0, Gen3 and Gen6), demonstrating unimpaired basal PSII photochemical performance in all the conditions and improved maximum photosynthetic electron transport capacity in Gen3. **e**, Heatmap of protein level in control and NP-treated samples quantifying protein abundance involved in carbon handling, RuBisCO and Calvin cycle, electron transport, photorespiration and TCA. Comparisons between treated and treated cultures (Gen0 vs Ctrl, Gen3 vs Ctrl, Gen6 vs Ctrl) and between early functionalized cells and high generation cells (Gen3 vs Gen0, Gen6 vs Gen0, Gen3 vs Gen6) were performed. Each box reports a color-coded log2 fold-change between the analysed and reference condition, for n=3 biological replicates. Red indicates the protein is upregulated compared to the reference condition; blue indicates the protein is downregulated compared to the reference condition; white indicates no change or protein not detected. Comparisons between conditions were performed and p-values after multiple hypothesis testing are reported in the heatmap as level of significance: * p<0.05, ** p<0.01, *** p<0.001.

Control cultures showed an initial increase in headspace CO_2_, reaching approximately 1500 ppm by day 4, followed by a gradual decline (Fig. 3b). Gen0 cultures accumulated substantially less CO_2_, reaching a maximum of approximately 750 ppm at day 6. In Gen3, CO_2_ peaked earlier, at days 4–5, and in Gen6 at days 3–4, both remaining at or below approximately 750 ppm before declining. Thus, NP functionalization consistently reduced net headspace CO_2_ accumulation potentially through changes in cellular respiration or CO_2_ consumption, NP-mediated carbon retention, or a combination of these processes. Notably, by Gen6 the temporal CO_2_ profile became more similar to that of untreated controls while remaining consistently lower in absolute concentration, suggesting progressive reorganization of carbon handling across successive subcultures.

Medium pH showed a stage-dependent response (Supplementary Fig. S11b,c). Gen0 cultures exhibited an initial alkaline shift followed by delayed acidification, whereas Gen3 and Gen6 progressively recovered more control-like dynamics. Although these changes broadly tracked headspace CO_2_ trends, extracellular pH is also influenced by ion, nutrient and metabolite exchange and therefore cannot be interpreted as a direct proxy for CO_2_ assimilation. Instead, the pH response is consistent with broader changes in extracellular metabolic regulation during successive NP functionalization (Supplementary Note 1).

We next examined whether altered CO_2_ dynamics were accompanied by changes in the photosynthetic apparatus. Chlorophyll accumulation increased progressively with successive subculture NP functionalization (Fig. 3c, Supplementary Table 1). Gen0 cultures showed a modest increase in chlorophyll content relative to untreated controls, reaching approximately 3% higher levels after 1 week and 26% after 2 weeks. The response was substantially greater in repeatedly functionalized cultures. After 1 week, chlorophyll content was approximately 120% and 160% higher in Gen3 and Gen6, respectively, compared with age-matched untreated controls, and remained elevated after 2 weeks of culture.

Pulse-amplitude-modulated (PAM) fluorescence was used to determine whether increased chlorophyll accumulation was accompanied by changes in photosynthetic performance (Fig. 3d and Supplementary Fig. S12-S13). The maximum quantum yield of photosystem II (Fv/Fm) showed modest reductions in Gen0 and Gen3 (-11% and -15%), consistent with an initial perturbation of photosystem II (PSII) function, whereas Gen6 recovered to control-like levels (-6% than control condition) (Fig. 3d-i). Gen0 also exhibited reduced maximum electron transport rate (ETRmax) (Fig. 3d-ii) and lower initial light-response slope (α) (-23% and -18% with respect to the control cultures, respectively), together with higher photoinhibition parameter (β) (+78% than control), (Supplementary Fig. S13a,b), indicating reduced photosynthetic performance following the initial NP functionalization.

In contrast, Gen3 displayed increased ETRmax (+46%) and lower β (-73%) compared to control, indicating an enhanced capacity to sustain electron transport across increasing irradiance with reduced high-light inhibition with respect to control cells. By Gen6, both ETR at saturation (ETRs) and ETRmax were constant compared with Gen3, while α recovered reaching control conditions, suggesting improved low-light efficiency and sustained maximum electron transport capacity between higher generations. PAM quantifications were performed on cultures exhibiting their characteristic aggregated phenotype, which may contribute by modifying light exposure within the cell population (Supplementary Fig. S12b, S13c).

However, the clear differences between the aggregated Gen3 and Gen6 suggested that the presence of aggregation alone is insufficient to explain the observed response and the photosynthetic response to repeated gPEI-Chi NP functionalization is stage dependent. Gen3 exhibits greater resistance to high-light exposure, while Gen6 reflects further reorganization of photosynthetic function, with higher low-light efficiency and maximum PSII quantum yield, rather than a broad upregulation of photosynthesis.

Untargeted proteomics supported this stage-dependent response (Fig. 3e, Supplementary Table 2, Fig. S14). In Gen0, several proteins associated with light harvesting and photosynthetic electron transport, including LHCB2.1, psbA, psbC, PSBO1 and PSAF, showed lower abundance relative to controls, together with proteins involved in Calvin–Benson cycle regeneration, including SBPase, TKL-1 and RPI3. Combined with the PAM measurements and the reduced representation of photosynthesis-related processes identified by gene set enrichment analysis a (GSEA; Supplementary Fig. S15), these changes are consistent with an initial perturbation of photosynthetic function following NP exposure.

In Gen3 and Gen6, proteins associated with light harvesting and energy production (LHCB2.1, RFNR1 and atpB), Calvin–Benson cycle regeneration (SBPase, TKL-1, RPI3 and TIM), and RuBisCO abundance and activation (rbcL, RBCS-1A and RCA) generally recovered relative to Gen0. These changes were consistent with the progressive development of starch-enriched chloroplasts observed by ultrastructural analysis. Enrichment of proteins associated with responses to high light was also consistent with the reduced photoinhibition observed following repeated functionalization. Recovery of LHCB2.1 coincided with the pronounced increase in chlorophyll content in Gen3 and Gen6 and may reflect changes in the abundance or organization of chlorophyll-containing antenna complexes.

However, because several PSII core proteins remained below control levels, the response is better characterized as a selective reorganization of the photosynthetic machinery rather than a uniform upregulation of photosynthesis. Changes in photorespiratory proteins, together with increased abundance of mitochondrial proteins, further indicate coordinated remodelling of photosynthetic carbon metabolism, photorespiratory recycling and mitochondrial energy metabolism in Gen3 and Gen6.

Carbonic anhydrases (CAs) abundance provided further evidence for adaptation of intracellular inorganic carbon handling. Whereas Gen0 showed relatively limited changes in CA abundance, Gen3 and Gen6 displayed a reorganization of the plastidial β-CAs. BCA1 and BCA2 were reduced in Gen3 and partially recovered in Gen6, whereas BCA5 increased significantly in Gen6. Together with the preserved PSII competence, reduced photoinhibition and persistently lower headspace CO_2_ concentrations, these changes are consistent with altered regulation of intracellular CO_2_/HCO₃⁻ pools during repeated NP functionalization. Importantly, RuBisCO subunits rbcL and RBCS-1A remained broadly comparable to controls in Gen3 and Gen6, indicating that the long-term response is not explained simply by increased RuBisCO abundance. Instead, the combined physiological and proteomic data point to broader reorganization of inorganic carbon handling, RuBP regeneration, photosynthetic energy production and carbon recycling.

Overall, gPEI–Chi NP functionalization over successive subcultures consistently reduced net headspace CO_2_ accumulation and progressively increased chlorophyll content, while inducing a stage-dependent reorganization of photosynthetic function. Gen3 exhibited the greatest photosynthetic electron transport capacity, whereas Gen6 combined recovered PSII quantum efficiency and sustained maximal electron transport. Together with the accompanying proteomic changes, these findings indicate that sustained NP functionalization progressively reorganizes carbon handling and photosynthetic metabolism rather than uniformly enhancing a single carbon-fixation pathway.

### NP functionalization alters cell wall regeneration dynamics

The plant cell wall is a major sink for photosynthetically derived carbon. We therefore investigated whether gPEI–Chi NP functionalization across successive subcultures altered cell wall regeneration using protoplasts as a single-cell model^39^. Control and NP-treated cultures were enzymatically digested after 4 days of growth, corresponding to active proliferation and high NP internalization, and transferred to cell wall regeneration medium. Viability, chlorophyll content and cell wall regeneration were monitored over 72 h (Fig. 4).

**Fig. 4.**
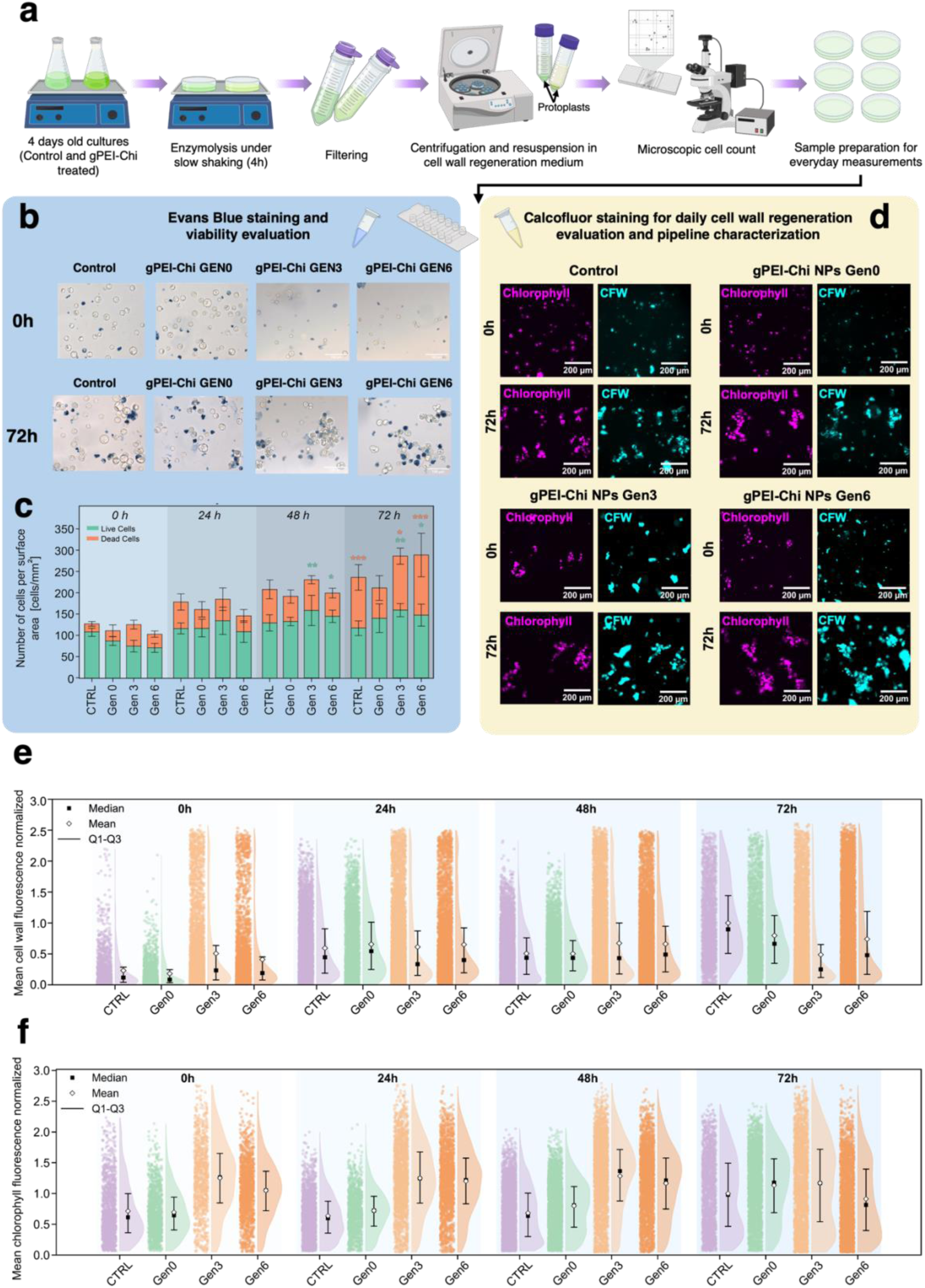
NPs functionalization alters cell wall regeneration dynamics in dedifferentiated plant cells. a, Schematic representation of the protoplasting workflow to generate dedifferentiated cell wall-free protoplasts from suspension cultures (created in BioRender.com). **b,** Protoplasts viability was tracked over three days of culture based on Evans blue staining for control and Gen0 cells and multi-generational NP-treated cell cultures (Gen3 and Gen6). **c,** Quantification on the brightfield images showed that NP functionalization does not impair short-term viability. **d,** An experimental pipeline for cell wall regeneration and chlorophyll analysis used chlorophyll autofluorescence and Calcofluor White (CFW) staining to image protoplasts for 3 days in control, Gen0, Gen3 and Gen6 protoplasts to track, respectively, cell wall deposition and chlorophyll dynamics in regenerating protoplasts. In the pipeline, from fluorescence images, cell wall **(e)** and chlorophyll **(f)** mean intensity of fluorescence were quantified per protoplast among the 3-days culture period and represented as scatter and violin plots, highlighting the median (filled square), mean (empty diamond) and first and third quartiles (Q1 and Q3).

Control and NP-treated protoplasts showed comparable viability immediately after isolation and subsequently underwent morphological recovery and cell division (Fig. 4b,c and Supplementary Figs. S16-S17). Viable cell density increased over time across conditions and remained predominant despite an increase in dead cells at 72 h. No significant differences in viability were detected between control and NP-treated groups at individual time points, indicating that NP functionalization did not compromise protoplast viability or regenerative capacity (Supplementary Table 3-6, Note 2).

Cell wall-associated fluorescence progressively increased in all conditions, but with distinct regeneration dynamics (Fig. 4d,e and Supplementary Figs. S18–S22; Supplementary Table 7-8, Note 3). Immediately after isolation, control and Gen0 protoplasts showed minimal wall-associated fluorescence, whereas Gen3 and Gen6 retained higher and more heterogeneous signals, consistent with less efficient digestion of the more aggregated cultures. During regeneration, control and Gen0 showed pronounced increases in wall-associated fluorescence and wall coverage, whereas changes were smaller and delayed in Gen3 and Gen6. Broad fluorescence distributions across conditions further revealed substantial cell-to-cell heterogeneity in wall regeneration. Together, these results indicate altered and slower wall regeneration in Gen3 and Gen6 cells.

Chlorophyll fluorescence also differed between conditions (Fig. 4d,f). Gen3 and Gen6 protoplasts initially exhibited higher and more homogeneous chlorophyll autofluorescence, consistent with their higher chlorophyll content before isolation, and remained comparatively stable during regeneration. In contrast, control and Gen0 populations progressively shifted towards higher chlorophyll fluorescence. Intracellular NP-associated fluorescence declined during regeneration, particularly in Gen3 and Gen6, while protoplast area remained comparable across conditions (Supplementary Fig. S23).

Overall, these single-cell analyses demonstrate that gPEI–Chi NP functionalization preserves protoplast viability and regenerative capacity while altering the kinetics and heterogeneity of cell wall regeneration. These distinct regeneration trajectories suggest that functionalization across successive subcultures establishes a cellular state with altered cell wall regeneration processes. We therefore investigated whether these changes were also reflected in the composition and organization of mature cell walls.

### NP functionalization progressively remodels cell wall architecture

Monosaccharide profiling revealed progressive changes in cell wall composition following NP functionalization (Fig. 5a and Supplementary Fig. S24). After one week, Gen0 cultures showed increased arabinose (+65.3%) and decreased rhamnose (−16.2%) and galacturonic acid (−6.5%), indicating an early shift in pectin composition. After two weeks, increased arabinose (+36%) and decreased galacturonic acid (−9.3%) persisted, accompanied by increased xylose (+19.3%). Gen3 and Gen6 similarly showed increased arabinose and xylose and decreased galactose relative to age-matched controls, while rhamnose and galacturonic acid remained largely unchanged. Glucose was also reduced in NP-treated cultures, particularly in Gen3 (−20.9%) and Gen6 (−17.2%), indicating broader remodelling of wall polysaccharide composition.

**Fig. 5.**
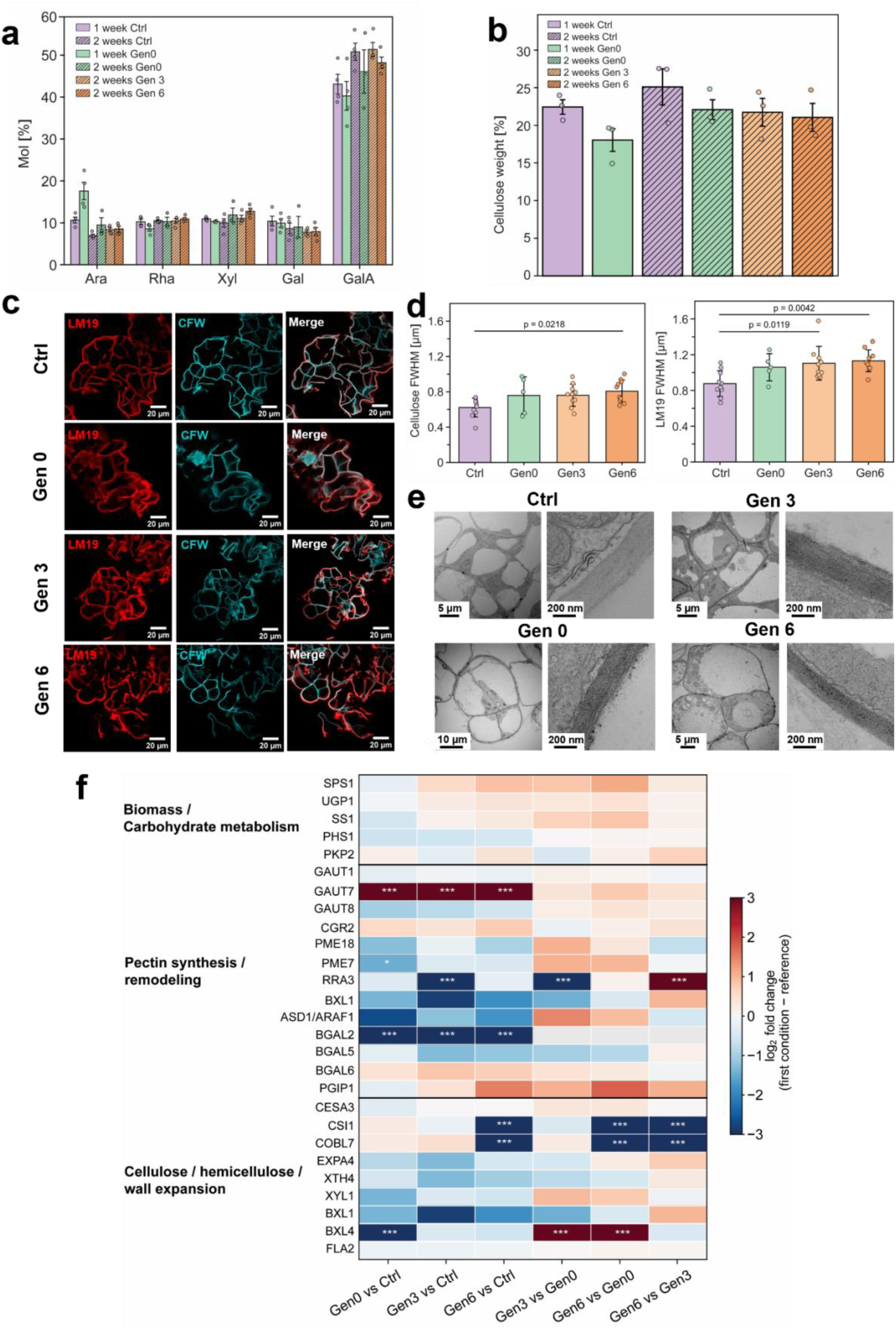
NP functionalization changes cell wall composition and architecture. **a**, Monosaccharide composition of extracted cell wall from control, Gen0, Gen3 and Gen6 cultures represented as mean ± standard deviation of moles percentage (n=3). Ara, Rha, Xyl, Gal and GalA represent arabinose, rhamnose, xylose, galactose, galacturonic acid, respectively. **b,** Cellulose content was evaluated as the percentage of mg of cell wall dry weight and represented as mean ± standard deviation (n=3). **c,** Immunofluorescence confocal images were acquired from cryo-thin sections of cells treated and non-treated with NPs, followed by immunostaining with monoclonal antibodies associated to low methylesterified pectin homogalacturonan (LM19) and Calcofluor White (CFW) for cellulose labelling in the cell wall. **d,** Full width at half maximum (FWHM) was quantified for cellulose and pectin fluorescence signal in the cell wall at the cell-cell interface to evaluate the effect of NP functionalization on cell wall thickness. **e,** Transmission electron microscopy images of control, Gen0, Gen3 and Gen6 cells showing ultrastructural changes both at the level of intracellular starch and chloroplasts and cell wall contrast. **f,** Heatmap of protein level in control and NP-treated samples quantifying protein abundance involved in biomass production and carbohydrate metabolism, pectin synthesis and remodelling, cellulose, hemicellulose and wall expansion. Comparisons between treated and treated cultures (Gen0 vs Ctrl, Gen3 vs Ctrl, Gen6 vs Ctrl) and between early functionalized cells and high generation cells (Gen3 vs Gen0, Gen6 vs Gen0, Gen3 vs Gen6) were performed. Each box reports a color-coded log2 fold-change between the analysed and reference condition. Red indicates proteins upregulated in the analysed condition; blue indicates proteins upregulated in the reference condition; white indicates no change or protein not detected. Comparisons between conditions were performed and p-values after multiple hypothesis testing are reported in the heatmap as level of significance: * p<0.05, ** p<0.01, *** p<0.001.

Cellulose quantification supported these compositional changes (Fig. 5b). Total cellulose remained relatively stable, with a modest reduction in Gen0 after one week and a gradual decrease at later stages, particularly in Gen6. Calcofluor White imaging further revealed transient cellulose-rich puncta following NP addition, persisting until day 3 in Gen0 and day 1 in Gen3 and Gen6, but absent from controls (Supplementary Figs. S25, S26), indicating localized reorganization of cellulose-rich wall domains.

To further investigate pectin organization, homogalacturonan (HG) immunolabeling revealed predominantly low- or non-methylesterified HG (LM19), with negligible highly methylesterified HG (LM20) across conditions (Fig. 5c and Supplementary Fig. S27). Although LM19- and cellulose-positive regions showed no major differences at cell– extracellular interfaces, both domains became broader at cell–cell interfaces in NP-treated cultures (Fig. 5d and Supplementary Fig. S28, Table 9). In Gen3 and Gen6, LM19-positive regions increased in apparent width by 26% and 29.1%, respectively, accompanied by increases of 22.3% and 29.8% in cellulose-positive domains. Thus, rather than substantially altering HG methylesterification, NP functionalization reorganized the spatial distribution of pectin- and cellulose-rich domains, particularly at cell–cell contact sites. The abundance of free carboxyl groups in low-methylesterified HG may also favour interactions with cationic gPEI–Chi NPs and contribute to their initial association with the cell wall^40,41^.

Cell wall ultrastructure provided further evidence of remodelling (Fig. 5e, Supplementary Fig. S29). Control cultures displayed smooth cell walls with homogeneous contrast and regular thickness. Gen0 walls retained a smooth appearance and regular thickness but showed increased contrast, whereas Gen3 and Gen6 developed a more granular ultrastructure. These changes may result from altered reorganization of cell wall fibre arrangement together with increased material accumulation, or changes in composition that enhance heavy metal staining.

Proteomic analysis supported extensive remodelling of cell wall biosynthesis and turnover (Fig. 5f and Supplementary Table 10). Gen3 and Gen6 showed increased abundance of proteins involved in sucrose metabolism and UDP-glucose production (SPS1 and UGP1), together with changes in starch metabolism (SS1 and PHS1), consistent with broader reorganization of carbon metabolism. Proteins involved in pectin biosynthesis and remodelling were also altered. Changes in GAUT proteins indicated reorganization of HG synthesis, while reduced abundance of RG-I-remodelling enzymes, including BXL1, ARAF1 and BGAL2/5/6, suggested decreased side-chain turnover. Together with the monosaccharide profiles, these changes support progressive reorganization of pectic domains rather than a general increase in pectin synthesis.

Cellulose-associated CESA3 showed only limited changes, consistent with the absence of a major increase in total cellulose. In contrast, proteins involved in xyloglucan remodelling (XTH4, XYL1), wall loosening (EXPA4) and polysaccharide rearrangement (BXL4), were reduced in NP-treated cultures. These changes potentially favour a denser and less dynamic architecture through stronger interactions between cellulose microfibrils and matrix polysaccharides, consistently with the increased contrast and granular wall ultrastructure observed in Gen3 and Gen6.

Changes in cytoskeletal organization and intracellular trafficking may further contribute to the altered wall regeneration observed in these cultures (Fig. 4d,e). Microtubule-associated proteins involved in nucleation and stabilization (TUBG1, TUBG2 and SPR1) increased without corresponding increases in proteins required for higher-order microtubule organization (AIR9 and MOR1), while myosin-associated motor activity decreased in Gen3 and Gen6 (Supplementary Fig. S30 and Table 11), potentially affecting the spatial organization and delivery of wall components during regeneration.

Together, these results demonstrate that gPEI–Chi NP functionalization progressively remodels cell wall composition and architecture. Changes in pectin biosynthesis and turnover, xyloglucan remodelling and wall-loosening processes are accompanied by broader pectin- and cellulose-rich domains at cell–cell interfaces and increasingly granular wall ultrastructure. This progressive wall remodelling provides a structural basis for the altered regeneration dynamics and increased cohesiveness of NP-functionalized cell aggregates.

## Conclusion

Here, we show that synthetic nanomaterials can be used to progressively engineer proliferating plant cell populations across successive subcultures. gPEI–Chi NPs were efficiently internalized by photosynthetic Arabidopsis suspension cells without compromising viability or proliferation. Supplying fresh NPs as the cultures were propagated enabled newly generated cells to be successively functionalized, leading to progressive changes in cellular and multicellular phenotype that were not apparent after the initial functionalization cycle.

This process extended beyond NP uptake. Successive functionalization reorganized carbon handling and photosynthetic physiology, producing persistently lower net headspace CO_2_ accumulation, increased chlorophyll content and stage-dependent changes in photosynthetic electron transport and associated metabolic pathways. In parallel, NP functionalization altered cell wall regeneration and progressively remodelled mature wall composition, spatial organization and ultrastructure. Broader pectin- and cellulose-rich domains at cell–cell interfaces, together with changes in wall biosynthesis and remodelling pathways, provide a structural basis for the increasingly cohesive multicellular aggregates that emerged across successive subcultures. Thus, the response to the NPs develops across multiple levels of biological organization, linking metabolic and photosynthetic changes with remodelling of the extracellular architecture.

These findings establish proliferating plant cell cultures as a platform in which nanomaterial functionality can be introduced as new cells are generated, allowing nano–bio interactions to shape the evolving phenotype of a living population. More broadly, this approach provides a route to engineer plant cells beyond transient modulation of individual physiological processes, coupling synthetic functionality to the intrinsic capacity of living cells to proliferate, remodel their metabolism and construct their own structural environment. Such population-level nanobionic engineering could provide a foundation for plant biohybrid systems and engineered living materials in which cellular physiology and material organization develop together over time.

## Materials and Methods

### Preparation of gPEI-Chi nanoparticles

The full synthesis procedure for gPEI-Chi nanoparticles and their characterization can be found in the supporting information of our previous study^26^.

### Cell culture preparation and subculture

All culture work and manipulations were conducted in sterile conditions. Calli for the preparation of stable cell lines and cell suspension cultures were obtained from root explants of Arabidopsis thaliana wild-type (Col-0), following the procedure described by *Pesquet et al.*^42^. The calli were grown on Murashige & Skoog (MS) medium (4.4 g/L, Duchefa Biochemie B.V., Haarlem, Netherlands) supplemented with 0.8% (*w/v*) agar (Sigma Aldrich, St. Louis, MO, USA) under dark conditions at 22°C. Once sufficiently grown, calli pieces measuring approximately 0.5x0.5cm were transferred to 6-well plates containing 3 mL of MS medium with 3% (*w/v*) sucrose (Sigma Aldrich, St. Louis, MO, USA) adjusted to a pH of 5.7. The calli pieces were dispersed into smaller fragments using sterile tweezers to facilitate cell release into the suspension medium. Sealed well plates were then placed back in the incubator (Pol-Eko aparatura, Model ST 700 CM, Poland) under dark conditions at 22°C with constant rotational agitation at 120 rpm. The plates were monitored daily, and once cell density was sufficiently high, 3 mL of cell suspension was transferred to 100mL Erlenmeyer flasks containing 27 mL of 3 % (*w/v*) sucrose MS medium. The flasks were sealed with cotton and aluminium foil and incubated in the dark at 22°C under continuous shaking at 120 rpm. After an adaptation period, 8 mL of cell culture was transferred to larger 250mL Erlenmeyer flasks containing 72 mL of MS medium with 3 % (*w/v*) sucrose. These flasks were again sealed with cotton and aluminium foil and maintained in the dark at 22°C while shaking at 130 rpm. Liquid cultures were subcultured weekly by directly performing a 1:10 dilution of cells in fresh culture medium (MS, 3 % (*w/v*) sucrose, pH 5.7) to promote continuous cell proliferation and maintain culture health.

Photosynthetic cell cultures were generated starting by established cell cultures kept in dark conditions. 8 mL of cells cultured in dark incubator were transferred in a 250 mL Erlenmeyer flask supplemented with 72 mL of MS medium with 1% (w/v) sucrose (pH 5.7) and then kept under continuous exposure to white light (150 μmol photons m^-2^ s^-1^) and constant shaking at 130 rpm and 22°C, with the final aim to induce the development of chloroplasts, following an established protocol from Dubreuil et al.^33^. The liquid cultures were subcultured weekly, by replacing the medium with fresh MS 1% sucrose first and then transferring the photosynthetic cells in the new flask with a 1:10 dilution in fresh MS 1% sucrose.

### Nanoparticle incubation in the cell culture

The gPEI-Chi nanoparticles were dissolved in sterile MS medium supplemented with 3 % (*w/v*) sucrose for cells in dark conditions and 1% (w/v) sucrose for cells under light conditions, at a concentration of 0.3 mg/mL and agitated until completely dissolved, following by a filtering step with a 0.22 µm sterile filter for ensuring sterile conditions. During the weekly subculture step, the cell culture medium was replaced with fresh medium and gently mixed. In a new flask, the 0.3 mg/mL gPEI-Chi NPs solution was diluted in fresh medium at a final concentration of 0.1 mg/mL in a total volume of 72 mL. Therefore, 8 mL of refreshed cell suspension was transferred to the new flask and gently mixed. The flask was sealed with cotton and aluminium and returned to the dark or light incubator under shaking and at 22°C. Cells were then monitored daily and utilized for all subsequent experiments as needed. The cells treated in the previously described way are reported as “Gen0” cultures, indicating a generation of cells exposed for the first time to the NPs.

### Multi-generational NPs-treated cultures

To underscore the long-term effect of continuous NP exposure on photosynthetic cell cultures, cells were subcultured by weekly adding fresh NP solution. In particular, starting with the Gen0 flask and every 7 days, the cell culture medium was replaced with fresh medium supplemented with NPs at a final concentration of 0.1 mg/mL and gently mixing them. 8 mL of cell suspension supplemented with NPs was transferred in a new flask keeping the regular 1:10 ratio in fresh medium supplemented with 0.1 mg/mL of NPs. This process was repeated for several weeks, reaching namely the “Gen3” after 3 weeks and the “Gen6” after 6 weeks, used as middle and final generations, respectively, for subsequent characterization experiments for long-term NP exposure.

### Growth curve measurements and cell volume after sedimentation (CVS)

To assess cell volume after sedimentation, a custom plastic vial holder was fabricated using a 3D printer (Ultimaker, model 2+ Extended, Utrecht, Netherlands). This holder was specifically designed to accommodate Erlenmeyer flasks containing the cultures, allowing them to be tilted at a 60° angle following a previously reported method^43,44^. After tilting, the flasks were held in position for five minutes to facilitate sedimentation of the cells. A ruler integrated into the apparatus enabled measurement of the height of the cell mass along the glass wall of the flasks and the cell volume was determined by referencing a calibration curve established using known volumes of water and their corresponding heights measured on the glass wall for flasks of similar volumes and tilt angles. The resulting volume of the cells, defined as the cell volume after sedimentation (CVS), was then expressed per litre of suspension.

### Dry weight measurement

Paper filter discs with a porosity of 100 mm were previously dried in oven at 60°C to remove the accumulated humidity and then weighed. 1 mL of cells was collected by each flask and filtered on the pre-weighed paper filter disc by using a Büchner vacuum funnel. The filtered cells on the paper disc were transferred in oven at 60°C for 24 h. The filter with dried cells was then weighed, registering the stable weight. The dry weight of the cells was calculated by subtracting the weight of the filter paper to the weight of cells plus filter.

### Evans Blue staining and viability

Evans Blue staining was carried out by first transferring 1mL of cell culture (with or without gPEI-Chi nanoparticles) into a small 2mL vial. Evans blue dye (DirectBlue 53, Sigma Aldrich, St. Louis, MO, USA) stock solution was then added to give a final concentration of 0.025% (*w*/*v*). The cells were stained at room temperature for 5 min before immediately being mounted for microscopy for further analysis. In light microscopy the living cells then appear unstained while dead cells appear blue stained.

### Cell wall Calcofluor White staining

Cell wall of plant cells was staining by using Calcofluor White stain. 1 mL of cell suspension was incubated with the staining solution at a concentration of 2.5 mM for 10 min before mounting for confocal microscopy inspection.

### Confocal Microscopy Imaging

Utilizing an inverted Zeiss LSM 980 confocal microscope equipped with an Airyscan2 detection unit, images were captured and processed using Zen Blue software version 3.4. Micrographs were acquired with a EC Plan-Neofluar 5x/0.16 M27, Plan-Apochromat 20x/0.8 M27 and Plan-Apochromat 63x/1.4 Oil DIC M27 objectives. Fluoresceine isothiocyanate (FITC) from the gPEI-Chi nanoparticles was excited with a 488 nm laser and detected using the AG520 filter (emission wavelength range 517-534 nm). Chlorophyll autofluorescence was acquired by exciting the sample with a 639 nm laser and collecting the signal by using a Chlorophyll A filter in the emission wavelength range 667-694 nm. Evans Blue fluorescence was detected by exciting it with a 639 nm laser and detected using the Alexa Fluor 680 filter (emission wavelength 702 nm). Calcofluor White stained cell wall was acquired exciting at 405 nm and collecting by using the filter CFW in the wavelength range 410-481 nm. LM19 and LM20 expression in the cell wall was detected by exciting with a 639 nm laser and collecting the emission within the range 650-700 nm.

### Absorption and fluorescence spectroscopy for NPs internalization

Absorption spectra for culture medium both with and without gPEI-Chi NPs were recorded on a microplate reader Biotek Synergy H1 in the range 300-700 nm with a step of 3 nm by using a Quartz 96 well plate. Fluorescence endpoint of culture medium with and without NPs was recorded at excitation 485 nm and emission 520 nm with gain fixed to 70.

### CO_2_ and pH measurements

A closed-loop CO_2_ measurement system was built to directly deliver the CO_2_ contained in the cell culture flask headspace to a CO_2_ sensor (Sensirion I2C SCD30) and an Arduino microcontroller board, pumped with a diaphragm gas micropump (NMP05KPDC-B 6V, Knf), as represented in Fig. 3a. In particular, the flask was sealed and connected to a water trap by means of impermeable tubes, after which a hydrophobic filter was used to maintain sterile condition during the measurement period. The diaphragm pump delivered the headspace CO_2_ from the flask to the sensor, which was sealed in a hermetic Petri dish to ensure any gas permeation during the measurement. CO_2_ was recycled back to the flask to measure the CO_2_ consumption/release during the observation time. The recording lasted 7 days and a registration step of 5 min was used for all the measured cultures.

Along with CO_2_ detection, pH was also measured every day for one week by using a Thermo Scientific Orion Star A211 pH Benchtop Meter. 20 mL cell culture medium was removed every day from the flask, filtered and used for measurement. A different flask was used every day to avoid altering the cell culture condition and the CO_2_-pH balance during handling and medium removal.

### Chlorophyll extraction and measurements

Chlorophyll was extracted from the control and NP-treated photosynthetic cell cultures at the seventh and fourteenth day of culture. For the 2-weeks conditions, after one week of culture, cell medium was refreshed without subculturing, keeping the culture for another week until the chlorophyll extraction. The extraction protocol was readapted from an optimized procedure^45^. The cell suspension was collected from the flask and dried with a paper filter disc by using a Büchner vacuum funnel. 500 mg of dry sample was weighed out for the extraction and transferred to a pestle, previously cooled down with liquid nitrogen. 260 mg of magnesium sulphate heptahydrate was added to the cells in the pestle, along with few droplets of liquid nitrogen and 15 mL of acetone. The mixture was grinded until complete homogenization and transferred and left for 15 min in a sealed beaker, after which it was filtered with a strainer with 20 μm mesh size and left to settle for other 10 min. The solution was centrifuged at 200 g for 5 min and the supernatant was used for absorbance measurement. A microplate reader Biotek Synergy H1 was used to record the absorbance spectra of the chlorophyll extracts between 350-700 nm with 3-nm step by using a Quartz 96 well. Absorbance at 647 nm and 665 nm, respectively was also collected and used for the chlorophyll concentration calculation. Chlorophyll a and b concentrations were calculated using the extinction coefficient reported by Inskeep et al^46^:

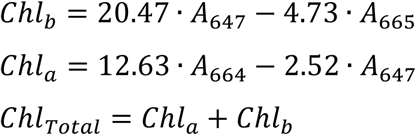

The final concentration of chlorophyll was computed as:

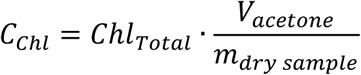

Where *V_acetone_* is the total volume of acetone used for the extraction and *m_dry_*_*sample*_ is the mass of the dried cell culture.

### Photosynthetic characterization

The photosynthetic performance of the samples was evaluated using a portable photosynthesis system (LI-6800, LI-COR Biosciences) equipped with a Multiphase Flash Fluorometer (6800-01A). The fluorometer was maintained in a horizontal position, and a custom 3D-printed black chamber was used to hold a Petri dish containing each liquid sample. Samples were dark-acclimated for 30 min before measurement. Illumination consisted of 90% red light (625 nm) and 10% blue light (475 nm).

Measurements were first performed under dark-adapted conditions to determine the minimum fluorescence (F_0_) and maximum fluorescence (F_m_). F_m_ was obtained by applying a rectangular saturating pulse with a duration of 1 s and an intensity of 6000 µmol photons m^-2^ s^-1^. Subsequently, the samples were exposed to increasing actinic light intensities of 10, 20, 50, 125, 150, 175, 300, 600, 900, 1200, and 1500 µmol photons m^-2^ s^-1^. At each light intensity, the steady-state fluorescence (F_s_) and the maximum fluorescence under illumination (F_m_^’^), obtained following application of the saturating pulse, were recorded.

From the fluorescence measurements, several photosynthetic parameters were calculated^47^. The maximum quantum yield of photosystem II (PSII), 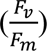, was calculated under dark-adapted conditions as:

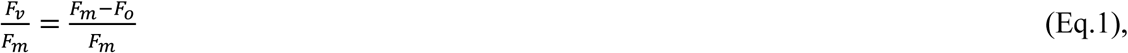

The effective quantum yield of PSII, (*φ_PSII_*), at each actinic light intensity was calculated as:

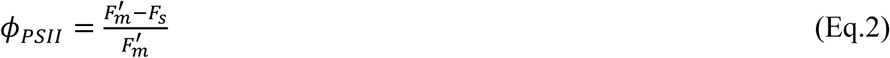

The electron transport rate (ETR) was subsequently estimated as:

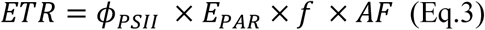

where (PAR) is the incident photon flux density integrated over the photosynthetically active radiation region (400–700 nm), *f* is the fraction of absorbed light allocated to PSII, assumed to be 0.5, and AF is the absorption factor, assumed to be 0.83. The latter should be considered an approximation, as the fraction of absorbed light can vary depending on parameters such as pigment content and cell density within the sample.

The resulting ETR vs light intensity curves were fitted using a custom python code and the model described by Platt and Gallegos (1980)^48^ :

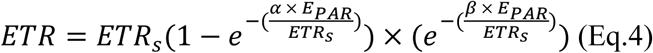

where *E_PAR_* is the incident light intensity, *ETR_s_* represents the theoretical maximum ETR in the absence of photoinhibition, α describes photosynthetic efficiency under low-light conditions and determines the initial slope of the ETR curve, and β describes the degree of photoinhibition at high light intensities. Thus, larger values of α indicate a stronger increase in ETR with increasing irradiance in the light-limited region, whereas larger values of β indicate stronger photoinhibition at high irradiance.

The maximum value of ETR was calculated after fitted as followed:

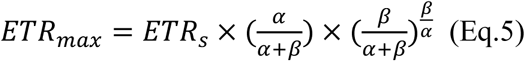

### Proteomic sample preparation

Photosynthetic cell cultures, treated and non-treated with gPEI-Chi NPs, were collected and pelleted, after which they were snap-frozen in liquid nitrogen. Pelleted cells were ground to a fine powder using a mortar and pestle in liquid nitrogen. The powdered pellet was thawed on ice and shortly equilibrated to room temperature. sodium deoxycholate (SDC) was added to 0.5 mL pellet to a final concentration of 3% (w/v) SDC. Cells were lysed using a FastPrep-5G homogenizer for six cycles of 45 s each. Samples were diluted to a final concentration of <1% SDC, and cell debris was removed by centrifugation at 21,000 × *g* for 5 min. Protein concentration was determined by Bradford assay with measured aliquots diluted to <0.1% SDC to avoid interference. For proteolytic digestion, 25 μg protein was reduced with 10 mM dithiothreitol (DTT) at 55 °C for 45 min and alkylated with 17 mM iodoacetamide (IAA) for 30 min in the dark. Proteins were then digested overnight (16 h) at 37 °C using a Trypsin/Lys-C Protease Mix (Thermo Scientific) at a protease-to-protein ratio of 1:100. Peptides were then desalted using StageTips packed with six layers of C18 Empore SPE disks (Merck) and dried in a SpeedVac at 45 °C for 1 h.

### Liquid chromatography–mass spectrometry (LC–MS) analysis

Samples were resuspended in 0.1% formic acid and analysed using a Q Exactive HF Hybrid Quadrupole-Orbitrap Mass Spectrometer operated in positive mode coupled to an UltiMate 3000 RSLCnano system equipped with an EASY-Spray ion source. Peptides were loaded onto a C18 Acclaim PepMap 100 trap column (75 μm × 2 cm, 3 μm particle size, 100 Å pore size) at a flow rate of 7 μL/min using solvent A (3% acetonitrile and 0.1% formic acid in water). Peptides were separated on an ES802 EASY-Spray PepMap RSLC C18 analytical column (75 μm × 25 cm, 2 μm, 100 Å) at 0.7 μL/min using a 40 min linear gradient from 1% to 32% solvent B (95% acetonitrile, 0.1% formic acid in water).

Peptides were analysed using data-dependent acquisition (DDA), with full MS1 spectra acquired at a resolution of 60,000 (at *m/z* 200) over an *m/z* range of 300–1200, followed by MS2 scans of the 10 most abundant precursor ions. MS2 spectra were acquired at a resolution of 60,000 using a 2 *m/z* isolation window and 20 s dynamic exclusion. The MS1 and MS2 maximum injection times were 205 and 105 ms, with AGC targets of 1 × 10⁶ and 2 × 10⁵, respectively. Precursor ions were fragmented by HCD at a normalised collision energy of 26.

### Proteomic data analysis

Raw MS data was analysed using MaxQuant (version 2.6.7.0). Spectra were searched against the *A. thaliana* FASTA reference proteome UP000006548 limited to reviewed entries. Trypsin/P digestion was used, allowing up to two missed cleavages, and match-between-runs enabled. Carbamidomethylation of cysteine was set as a fixed modification, while methionine oxidation and protein N-terminal acetylation were set as variable modifications. Identifications were filtered at a 1% false discovery rate (FDR). Proteins were median-normalized, and differential expression was calculated using MSstats (version 4.4.1) in R^49^. Gene set enrichment analysis (GSEA) was performed in R using the ClusterProfiler package (version 4.15.1)^50^.

### Protoplasts extraction

The protoplasts extraction protocol is an adaptation of the protocols from Yoo et al.^51^ and Bogdziewiez et al.^39^ and it is summarized below. A protoplasting buffer (PB) was prepared adding in deionized water 20 mM MES, 0.4 M mannitol, 20 mM KCl, 10 mM CaCl_2_, 0.1% (w/v) BSA and adjusting the pH to 5.7. An enzymatic solution was prepared for the cell wall lysis and protoplasts extraction by warming up the PB solution to 55°C under stirring and adding 1.5% (w/v) Cellulase R-10 and 0.4% (w/v) Macerozyme R-10. Under sterile conditions, 5 mL of 4-days-old control photosynthetic cell suspension and 15 mL of 4-days-old photosynthetic cells treated with gPEI-Chi NPs (Gen0, Gen3 and Gen6) were transferred and the medium was replaced with 20 mL of the filtered enzymatic solution. Different volumes of cell suspensions were used between treated and non-treated conditions, balanced over the differences in CVS detected at day 4 of culture (Fig. 1e-i). Cells were gently mixed with the digestion solution and 10 mL of the cell suspension was transferred to glass non-treated Petri dishes, followed by incubation in dark conditions for 4 hours under mild rotation (74 rpm) at 22°C. A high-salt solution (W5 solution) was prepared to inhibit the enzymatic reaction, by adding 2 mM MES, 154 mM NaCl, 125 mM CaCl_2_, 5 mM KCl in deionized water and adjusting its pH to 5.7. To stop the enzymatic digestion, the sterile W5 solution was added to the digested cell culture in a ratio 1:1 and incubated back for 5 min under mild rotation (30 rpm). The cell suspension was then filtered with a 70 µm cell strainer to isolate single protoplasts from bigger clusters of cells. To remove debris and digested cell wall material, the protoplasts were centrifuged at 200g for 5 min, the supernatant was removed and protoplasts were resuspended in a cell wall regeneration medium (CWR) made by 0.316% (w/v) Gamborg B5, 20 mM MES, 0.25 M mannitol, 0.2 M glucose, 5 µM NAA in deionized water and calibrated to achieve pH 5.7. Protoplasts were counted, seeded in non-treated plastic Petri dishes at a density of 2 · 10^5^ cells/mL in CWR and incubated at 22°C in dark conditions without shaking. Independent samples were used every day for viability evaluation through Evans Blue assay, cell wall regeneration detection with Calcofluor White staining and chlorophyll content quantification at 0, 24, 48 and 72 h.

### Protoplasts’ wall regeneration - Data analysis

Every day starting by the day of protoplasting (day 0), protoplasts were stained with Evans blue and loaded in prefabricated microchannel chips (IBIDI µ-Slide VI Flat, non-treated) for imaging. Protoplasts were acquired with a fluorescence microscope Nikon Eclipse Ni-E equipped with a 20x objective in brightfield with a color-coded camera. At least 30 images per microchannel were acquired for each condition. Dead cells were counted as blue-stained circular protoplasts, while live cells were considered as round-shaped, intact and non-stained protoplasts. The count of live and dead cells was normalized to the total acquired area in mm^2^.

In parallel, protoplasts were stained with Calcofluor White stain and loaded in a parallel microchannel. Multichannel images were acquired with the Nikon Eclipse fluorescence microscope with a 20x objective, acquiring consecutively brightfield, NP-associated fluorescence (FITC filter, 488 nm), chlorophyll autofluorescence (Cy5 filter, 550 nm) and cell wall-associated Calcofluor signal (DAPI filter, 370 nm). At least 20 images per microchannel were acquired for each condition. A custom Python code was used for collecting the multichannel images, creating 2-by-2 tiles of the separate 4 channels to create an hyperstack for successive data analysis. Following the Q-Warg pipeline procedure previously developed by Bogdziewiez et al.^39^, a mask was generated by segmentation of the protoplasts’ brightfield tiles by using CellPose, a deep learning-based ImageJ (Fiji) macro to automate objects segmentation. A cyto4 pretrained model developed by Stringer and Pachitariu^52^ was used to obtain the labelled mask for each segmented protoplast. The Q-Warg pipeline was followed by using an ImageJ macro extracting the morphometric parameters from the segmented labels, including cell size and circularity, as well as to measure the mean intensity of fluorescence related to chlorophyll, cell wall and gPEI-Chi NPs for each segmented protoplast, by using the MorphoLibJ library. A custom Python script was instead used for previous data plotting through an interactive GUI for manual selection of the parameters of interest for the analysis. In particular, the mean fluorescence intensity for each detected protoplast was matched to the corresponding morphometric and chlorophyll measurements using the label associated during segmentation. Protoplasts were initially filtered according to morphology and chlorophyll thresholding, retaining objects with a circularity greater than 0.88 and chlorophyll fluorescence intensity greater than 100, aimed to represent only protoplasts classified as alive when they retain the circular shape and possess chlorophyll. The data on cell wall and chlorophyll mean fluorescence intensity for each protoplast were then normalized to the corresponding averaged mean fluorescence of the protoplasts’ control population at day 3 for each experiment. Differently, NPs mean fluorescence intensity for each protoplast was normalized to NPs maximum fluorescence. These normalization steps were included to account for differences between experimental sets and represent the data as relative increase/decrease of fluorescence intensity with respect to maximum conditions of cell wall, chlorophyll content and NP-related fluorescence. All the data related to normalized cell wall mean fluorescence, normalized chlorophyll mean fluorescence, normalized NP mean fluorescence and protoplasts’ area were visualized using violin plots of the distribution of the single protoplasts mean fluorescence intensity and area for each experimental condition, underlying the mean, median and interquartile range of the 25^th^ and 75^th^ percentile (Q1-Q3).

Cell wall coverage progression during the protoplasts culture in CWR for the 4-day observation time was quantified using a custom ImageJ macro as the fraction of protoplast area positive for cell wall staining. First, the fluorescence images of protoplasts were background-subtracted and segmented following the CellPose procedure previously described. The background-corrected cell wall channel was converted into a binary mask by Otsu thresholding, classifying pixels as either cell wall-positive or background. For each segmented cell, the mean intensity of the binary mask was extracted and the percentage of the cell area occupied by cell wall fluorescence was computed as:

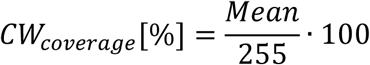

The mean ± standard deviation of cell wall coverage between the protoplasts’ population was calculated for each condition and time point.

### Cell wall extraction and monosaccharides quantification

Photosynthetic cell cultures (control, Gen0, Gen3 and Gen6) were freeze-dried and subsequently homogenized to a fine powder using a ball mill. Alcohol insoluble residue (AIR) was prepared from 35 mg of ground material per sample by first incubating it with 80% ethanol for 30 min at RT which was followed by 30 min incubation with chloroform: methanol (1:1). The resulting pellet was dried and treated with 1000 units of α-amylase in 0.1M potassium phosphate buffer to remove starch. After overnight incubation, the samples were successively washed with Milli-Q water and acetone then dried overnight in a vacuum desiccator.

For monosaccharide composition analysis, around 500 μg of dry weight (DW) of the processed AIR was derivatized via trimethylsilylation (TMS) and analysed by GC/MS (7890A/5975C; Agilent Technologies) according to previously described protocols^53^. The degree of methyl esterification was determined using around 1.5 mg DW of AIR. Samples were first saponified with 1 M NaOH for 1 h at RT, neutralized with 1 M HCl, and approximately 180 µL of the supernatant was collected for methanol quantification. Reactions were prepared in ELISA plates: 50 µL of alcohol oxidase in 0.1 M sodium phosphate buffer was added to each sample and incubated for 15 min at RT. Then, 100 µL of a developer solution (containing acetylacetone, acetic acid, and ammonium acetate in water) was added, plates were sealed and incubated for 10 min at 68°C. Absorbance was measured at 412 nm. The analysis of the absorbances was done as previously described^54^.

Crystalline cellulose was isolated using the previously described Updegraff method^55,56^. The obtained glucose was measured using the anthrone assay^57^.

Data was further analysed and the graphs were generated with the help of RStudio.

### Cryosectioning for cell wall composition

Photosynthetic plant cells, treated and non-treated with gPEI-Chi NPs, were prepared for cryosectioning and immunostaining, with the aim to characterize the cellulosic and pectinic contribution to the cell wall composition before and after NP functionalization. In more details, cells were collected after 1 week of culture in light conditions, washed 3 times in 1x PBS and fixed in 4% PFA for 30 minutes. Cells were then washed 4 times in PBS and embedded in gradient of sucrose for reducing the formation of ice crystals and protect cell structures from low temperatures damage. In particular, PBS was removed and cells were gradually incubated in 10% (w/v) sucrose for 25 min, 20% (w/v) sucrose for 45 min and overnight incubated at 4°C with 30% (w/v) sucrose. Sucrose was removed and cells were transferred to cryomolds (Simport Scientificinc M475), covered with optimal cutting temperature (OCT) compound (CellPath KMA-0100-00A) and snap-frozen in liquid nitrogen vapours for few minutes until OCT solidification was achieved. Samples were then stored at -80°C. Cryosections of 5-10 µm were obtained transferring the embedded cells on cryostat stabs and cutting them with cryostat microblades (MX35 ultra, Thermo Scientific) using a cryostat (Epredia CryoStar NX70) at -15°C. Sections were then collected onto adhesion slides (Epredia™ SuperFrost Plus™, 10149870, Fisher Scientific), air-dried for few hours and stored at 4°C.

### Immunofluorescence for cell wall composition

The sections were washed 3 times in 1x PBS and once in PBS-T (1x PBS and 0.1% Tween 20) for 5 min. They were then incubated in blocking buffer consisting of 1% (w/v) BSA in PBS-T for 1 h to prevent non-specific labelling. After 3 washes in PBS-T, the sections were incubated with the anti-pectic monoclonal primary antibodies LM19 and LM20 (Agrisera) diluted in 0.1% (w/v) BSA in PBS-T at a ratio of 1:20 for 2 h at room temperature in humid chamber and darkness. After 3 washes in PBS and 3 washes in PBS-T, the secondary antibody Alexa Fluor 647 (Invitrogen™ Goat anti-Rat IgM, Heavy chain, Fisher Scientific) diluted 1:100 in blocking buffer 0.1% (w/v) BSA in PBS-T for 2 h at room temperature or overnight at 4°C in humid chamber and darkness. After washing in PBS-T, the sections were also stained with Calcofluor White stain diluted at a concentration of 2.5 mM in PBS-T for cellulose imaging. An anti-fade medium made by mixing PBS and glycerol (1:1) was used, the sections were sealed and stored at 4°C in darkness until imaging.

### Data analysis for cell wall composition

Confocal images were acquired for the sectioned cells, acquiring multichannel images of LM19 and Calcofluor White fluorescence and LM20 and Calcofluor White fluorescence. Pectin- and cellulose-associated wall thickness was estimated from fluorescence intensity profiles acquired across LM19+LM20 and Calcofluor positive regions both at the boundaries of individual cells and at the cell-cell interface between adjacent cells. Prior to analysis, the images were processed in ImageJ by subtracting the background. Multiple line were manually drawn approximately perpendicular to the cell boundary and crossing the fluorescent cell wall signal, while the corresponding intensity profiles were extracted as grey-value intensity versus distance. The line profiles were analysed using a custom Python script. Here, for each line profile the fluorescence baseline (*I_baseline_*) was defined as the minimum intensity value while the peak (*I_peak_*) was defined as the maximum intensity. The signal amplitude was then calculated as the difference between the peak and baseline, retrieving the half-maximum intensity (*I_HM_*) as:

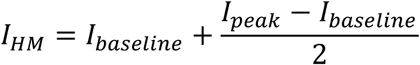

The values at which the fluorescence profile crossed the half maximum level on either left and right side of the peak were determined by linear interpolation. The full width at half maximum (FWHM) was then calculated as the distance between the left and right half maximum crossing points. This provided an estimation of the apparent width of the cellulose- and pectin-positive area in the cell wall of NP treated and non-treated photosynthetic cells. At least 15 lines were used for line profile detection for each cell (for boundary interface) or pair of cells (for cell-cell interface) and the resulting FWHM values were averaged to obtain a single mean value per cell. At least 10 cells were analysed per condition and the values were reported as mean ± standard deviation.

### Transmission electron microscopy (TEM)

Cells from 1-week control and NP-treated cultures were transferred and incubated with fixative solution made by 2.5% glutaraldehyde (EM grade) in 0.1 M phosphate buffer for 10 minutes. The solution was removed letting the cells settle and replaced with 2h postfixing 1% osmium tetraoxide (OsO_4_) in water. After that the samples were dehydrated in ethanol gradient followed by resin infiltration (3:1, 1:1, 1:3 resin-to-pure ethanol) and embedded in pure Spurr’s resin ( TAAB Laboratories Equipment Ltd, England). To prepare sections the samples in resin were trimmed, and the presence of the cells were confirmed under a light microscope. Ultra-thin sectioning of approximately 70 nm thickness was done using an Ultra microtome (Leica EM UC7) and the sections were placed on hexagonal copper grids, with a formvar film. The ultra-thin sections were stained with uranyl acetate (EMS England 22451) and lead citrate and examined with the Thermo Scientific Talos L120C transmission electron microscope.

### Statistical analysis

For the chlorophyll and photosynthetic characterization through pulse amplitude modulated (PAM) chlorophyll fluorescence, three technical replicated were measured for each biological replicate. Technical replicates were first averaged to obtain a mean value for each biological replicate. Normality and homogeneity of variance were tested by using Shapiro-Wilk and Levene’s test. One-way ANOVA followed by Tukey’s multiple comparison test were subsequently performed using the biological replicates as independent observations. Data are reported as the mean ± standard deviation among biological replicates. Significance levels were indicated as * p<0.05, ** p<0.01, *** p<0.001.

In the proteomic data analysis, proteins differential expression were statistically tested with p-values adjusted for multiple hypothesis testing using the Benjamini-Hochberg method. Proteins detected in fewer than n-1 replicates were excluded from statistical analysis. The log2 fold change (log2FC) values in the heatmaps were defined as the abundance in the first condition minus that in the reference condition, with positive and negative values indicating increased and decreased protein abundance, respectively. Significance levels were indicated as * p<0.05, ** p<0.01, *** p<0.001. In the gene set enrichment analysis (GSEA) proteins were ranked according to their t-statistic from the differential expression analysis and analysed using the GSEA function. Default settings were used (gene set size range of 10–500 and Benjamini– Hochberg correction for multiple testing) with a p-value cutoff of 0.05.

Protoplasts viability was assessed by analysing live and dead cell densities. Homogeneity of variance among groups was evaluated using Levene’s test. As the assumption of equal variances was satisfied, data were analysed using ordinary two-way ANOVA, with experimental condition and time as fixed factors, including their interaction. Planned post-hoc comparisons were performed to compare each NP-treated group with the corresponding control at individual time points and to evaluate temporal changes within each condition relative to day 0 of protoplasting. These multiple comparisons were corrected using the Holm method and differences were considered statistically significant at p<0.05 and indicates as * p<0.05, ** p<0.01, *** p<0.001.

For protoplasts cell wall and chlorophyll content evaluation, mean intensity of fluorescence distributions were non-normally distributed and non-parametric rank-based statistics were applied. Differences among multiple groups were assessed using the Kruskal-Wallis test, separately comparing conditions within each time point and then time points within each condition. When significant differences were detected, pairwise comparisons were performed using the Mann-Whitney U test followed by Holm correction for multiple comparisons. Statistical significance was defined as adjusted p<0.05 and indicates as * p<0.05, ** p<0.01, *** p<0.001.

Protoplasts’ cell wall coverage was analysed testing the mean values with a two-way ANOVA with condition and time as fixed factors and their interaction. Before, homogeneity of variance was assessed using Levene’s test. Planned post-hoc comparisons were performed among conditions at each time point and comparisons of each time point with day 0 within the same condition. P-values were adjusted using Holm correction for multiple testing. Statistical significance was defined as adjusted p<0.05 and indicates as * p<0.05, ** p<0.01, *** p<0.001. For the monosaccharides quantification analysis, statistically significant differences between the samples were determined using Kruskal-Wallis with Dunn’s test for pairwise comparisons (P<0.05) and Wilcoxon rank-sum test (P<0.05) where applicable. Statistical significance was defined as adjusted p<0.05 and indicates as * p<0.05, ** p<0.01, *** p<0.001.

Cellulose and pectin thickness quantification at single-cell level was tested by analysing the mean FWHM measured across cell boundary and cell-cell contact, by comparing control and NP-treated cell cultures. Data distribution within each condition was assessed using the Shapiro-Wilk test, while homogeneity of variance among groups was tested using Levene’s test. When normality and equal variance assumptions were satisfied, differences among conditions were analysed using one-way ANOVA followed by Tukey’s multiple comparison test. When parametric assumptions were not met, the Kruskal-Wallis test was used, followed by pairwise Mann-Whitney U test with Bonferroni correction for multiple comparisons. Statistical significance was defined as adjusted p<0.05 and indicates as * p<0.05, ** p<0.01, *** p<0.001.

## Acknowledgments

The authors wish to thank Agnieszka Ziolkowska (Umeå University) for TEM imaging and support for the discussion for ultrastructural characterization of functionalized cells.

The authors thank Therése Klingstedt (Department of Physics, Chemistry and Biology (IFM) – Linköping University) for support for cryostat sectioning of functionalized cells.

The work was primarily supported by the European Union (ERC-2021-STG, 4DPhytoHybrid, 101042148) and the Knut and Alice Wallenberg Foundation (KAW project FATE 2022.0029). Additional funding was provided by the Swedish Government Strategic Research Area in Materials Science on Advanced Functional Materials at Linköping University (Faculty Grant SFO-Mat-LiU No. 2009-00971).

The authors acknowledge the facilities and technical assistance of the Umeå Core Facility Electron Microscopy (UCEM) at the Chemical Biological Centre (KBC), Umeå University, a part of the National Microscopy Infrastructure NMI (VR-RFI 2019-00217).

This work was supported by grants from the Novo Nordisk Foundation (NNF21OC0067282) and Åforsk Foundation (20-502).

This work was also supported by Umeå Plant Science Center with grants from the Knut and Alice Wallenberg Foundation (KAW 2016.0352 and KAW 2020.0240); the Wallenberg Initiatives in Forest Research (WIFORCE) funded by the Knut and Alice Wallenberg Foundation, the Swedish Governmental Agency for Innovation Systems (VINNOVA 2016-00504); and Bio4Energy, a Strategic Research Environment supported through the Swedish Government’s Strategic Research Area initiative.

## Author contributions

E.S., C.R. and D.P. conceived and designed the research. E.S. supervised the work. C.R. established the NP-treated cell culture and protoplasts models, performed initial experiments in NP internalization, set the CO_2_ measurement apparatus and performed initial CO_2_ experiments, performed CVS and dry weight measurements on both non- and photosynthetic cells, chlorophyll extraction and measurement. D.P. performed the NP internalization experiments and characterization, the CO_2_ measurements and chlorophyll extraction and measurement, cell chlorophyll/wall and NP/chlorophyll colocalization imaging, pH measurements, protoplasts extraction and full characterization, LM19/20/cellulose immunofluorescence and analysis, proteomics data interpretation, and all the statistical analysis. A.K. and E.P.H. conducted the proteomics experiments and data analysis, providing support for the discussion. L.L. and S.V. performed the measurement and analysis on cell wall monosaccharides and cellulose content, and contributed to the discussion for cell wall composition. L.Boutonnet refined the pipeline for protoplasts cell wall and chlorophyll quantifications. L.Bogdziewiez and S.V. provided protocols and support for protoplasts extraction, maintenance and data analysis. X.J. and A.S. provided calli of root plant cells for starting the cell cultures and provided expertise in plant cell culture. E.C. established the protocol of transition from dark cell culture to light cell cultures and performed CVS and dry weight measurements of non-photosynthetic cells. A.G.F. performed the photosynthetic measurements and analysis. D.M. synthesized and provided the green fluorescent polyethyleneimine-chitosan nanoparticles (gPEI-Chi NPs) for all the experiments performed in the paper. E.S., D.P. and C.R. wrote the manuscript with input from all authors.

## Data availability

The data that support this study are all included in the manuscript and raw data are available upon request from the authors.

